# Benchmarking inverse statistical approaches for protein structure and design with exactly solvable models

**DOI:** 10.1101/028936

**Authors:** Hugo Jacquin, Amy Gilson, Eugene Shakhnovich, Simona Cocco, Rémi Monasson

**Affiliations:** Laboratoire de Physique Statistique de l’Ecole Normale Supérieure - UMR 8550, associé au CNRS et à l’Université Pierre et Marie Curie, 24 rue Lhomond, 75005 Paris, France; Department of Chemistry and Chemical Biology, Harvard University, 12 Oxford Street, Cambridge, MA 02138, USA; Laboratoire de Physique Théorique de l’Ecole Normale Supérieure - UMR 8549, associé au CNRS et à l’Université Pierre et Marie Curie, 24 rue Lhomond, 75005 Paris, France

## Abstract

Inverse statistical approaches to determine protein structure and function from Multiple Sequence Alignments (MSA) are emerging as powerful tools in computational biology. However the underlying assumptions of the relationship between the inferred effective Potts Hamiltonian and real protein structure and energetics remain untested so far. Here we use lattice protein model (LP) to benchmark those inverse statistical approaches. We build MSA of highly stable sequences in target LP structures, and infer the effective pairwise Potts Hamiltonians from those MSA. We find that inferred Potts Hamiltonians reproduce many important aspects of ‘true’ LP structures and energetics. Careful analysis reveals that effective pairwise couplings in inferred Potts Hamiltonians depend not only on the energetics of the native structure but also on competing folds; in particular, the coupling values reflect both positive design (stabilization of native conformation) and negative design (destabilization of competing folds). In addition to providing detailed structural information, the inferred Potts models used as protein Hamiltonian for design of new sequences are able to generate with high probability completely new sequences with the desired folds, which is not possible using independent-site models. Those are remarkable results as the effective LP Hamiltonians used to generate MSA are not simple pairwise models due to the competition between the folds. Our findings elucidate the reasons of the power of inverse approaches to the modelling of proteins from sequence data, and their limitations; we show, in particular, that their success crucially depend on the accurate inference of the Potts pairwise couplings.

**Author Summary:** Inverse statistical approaches, modeling pairwise correlations between amino acids in the sequences of similar proteins across many different organisms, can successfully extract protein structure (contact) information. Here, we benchmark those statistical approaches on exactly solvable models of proteins, folding on a 3D lattice, to assess the reasons underlying their success and their limitations. We show that the inferred parameters (effective pairwise interactions) of the statistical models have clear and quantitative interpretations in terms of positive (favoring the native fold) and negative (disfavoring competing folds) protein sequence design. New sequences randomly drawn from the statistical models are likely to fold into the native structures when effective pairwise interactions are accurately inferred, a performance which cannot be achieved with independent-site models.

## Introduction

Prediction of protein structure from sequence remains a major goal of computational structural biology with significant practical implications. While great progress was achieved in de novo prediction of structures and even direct folding of smaller globular water soluble proteins [1,2], structure prediction remains challenging for larger proteins and membrane and other non-globular proteins. For these cases indirect methods such as homology modeling are most promising. An approach to predict structure from the statistics of multiple sequence alignment (MSA) was proposed twenty years ago [3,4]. The underlying assumption for these statistical approaches is that residues that covary in a MSA are likely to be in close proximity in protein structure.

Recently, this approach has been significantly improved and has become a practical tool to extract structural information from sequence data [5,6], and to help folding proteins [7]. Progress was made possible by the application of the maximum entropy principle of statistical mechanics [8,9] to derive the distribution of sequences in a protein family, *i.e*. their probability to appear in a MSA. This approach is similar in spirit to the derivation of Gibbs distribution in Statistical Mechanics, with an effective Hamiltonian constructed to reproduce single-site amino-acid frequencies and pairwise amino-acid correlations. As such the effective Hamiltonian is a Potts model [10], whose parameters (the site-dependent fields and pairwise couplings acting on amino acids) are fitted to reproduce the MSA statistics. This approach allows one to disentangle direct (corresponding to couplings) from indirect (mediated by other sites) correlations [6, 11]; large couplings are much better predictors of contacts than large correlations.

The exploitation of covariation information in proteins extends beyond structural prediction, and is potentially useful for homology detection, for characterizing the fitness changes resulting from mutations, or for designing artificial proteins with ‘natural’ properties [12]. Experiments show in particular that a sizeable fraction of artificial sequences generated to respect the 1- and 2-point amino-acid frequencies calculated from the natural MSA of the WW domain (a short protein domain with ⋍30 amino acids) acquire the native fold [13, 14]. This fraction vanishes when artificial sequences are generated which respect the pattern of single-site frequencies only. Recent studies have shown that inferred maximum-entropy models are helpful to predict the effects of mutations in HIV virus sequences [15].

Despite those successes fundamental issues remain poorly understood. To what extent do the inferred couplings reflect the physical (energetics) and structural (contact) properties of proteins? How much are the covariation properties of one family influenced by the need to prevent sequences from folding in competing structures? Are pairwise correlations, and hence, couplings generally sufficient to capture the statistical features of protein families and define generative models?

In this work we answer those questions using lattice-based protein (LP) models [16-18]. Using LP allows us to evaluate the strengths and limitations of the maximum-entropy Potts modelling and of the inference methods to approximate the Potts pairwise couplings by comparing recovered solutions with known exact ones. Protein-like sequences are generated using a well-defined Hamiltonian that employs 20 amino-acid types and Miyazawa-Jernigan energy function in contact approximation [19]. LP share many common properties with real proteins, for instance non trivial statistical features of the sequences associated to a given fold, with variable conservation along the protein chain [17, 20]. Surprisingly, the covariation properties of LP had not been much studied so far [21]. Here, we do so by generating MSA data for various LP folds, and apply to those sequence data the inverse approaches used for real proteins (Fig. 1). In particular, we show that the inferred couplings are excellent predictors of contacts in the protein structure. In addition, while the inferred Potts pairwise couplings mostly reflect the energetics and the contacts in the fold corresponding to the MSA, they also strongly depend on the nature of the other folds competing with the native structure, and have transparent interpretations in terms of positive and negative designs. Furthermore, we show that the pairwise Potts model (when carefully inferred) is generative: it produces with high probability sequences with the right fold, a performance which cannot be achieved by models reproducing single-site frequencies alone. This is a non-trivial result as the log probability of LP sequences constrained to fold in a given structure is not a sum of pairwise contributions only, but includes multi-body interactions to all orders due to contributions from competing folds.

**Figure 1:**
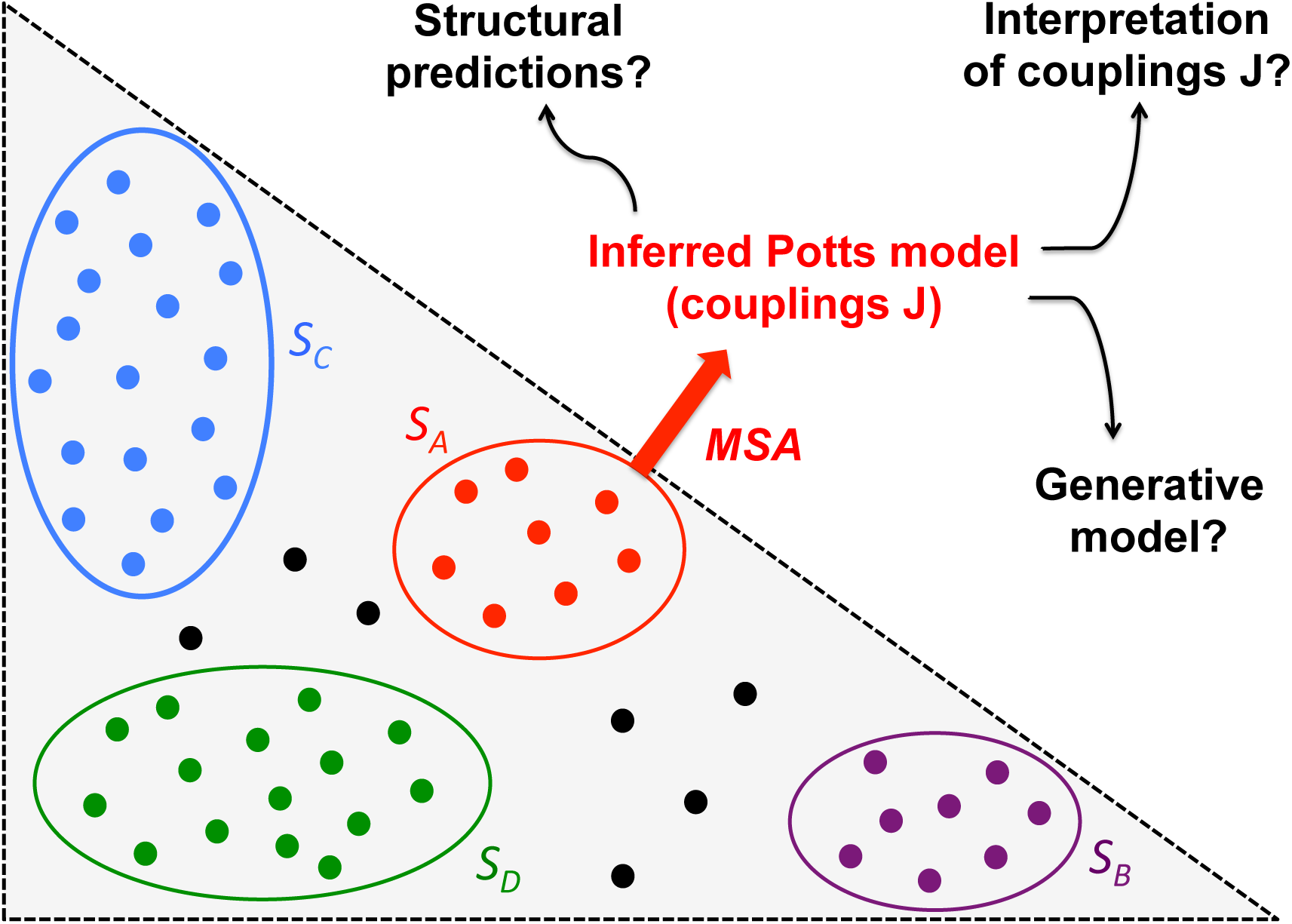
From lattice-protein sequence space to inferred Potts model. Protein families, each corresponding to a particular structure *S*, represent portions of sequence space (colored blobs), in which all sequences (colored dots) fold on a unique conformation. Many sequences are expected to be non folding, and not to belong to any family (black dots). Protein families differ by how much they are designable, *i.e.* by the numbers of sequences folding onto their corresponding structures, represented here by the sizes of the blobs. *S_A_* and *S_B_* are the least designable folds, while *S_C_* and *S_D_* are realized by larger numbers of sequences, see SI Table. I. From a multi-sequence alignment (MSA) of one family, we infer the maximum-entropy pairwise Potts model reproducing the low-order statistics of the MSA. The model is then tested for structural prediction and generating new sequences with the same fold. An important issue is to unveil the meaning of the inferred pairwise couplings **J**, which depend both on the family fold, as well as on the competitor folds.

## Results

We have generated MSA for four representative LP structures shown in Fig. 2, referred to as *S_A_* to *S_D_*; the energetics of the LP model, the Boltzmann probability *P_nat_*(*S*|**A**) of a sequence **A** to achieve a fold *S*, and the procedure for sampling the sequence space and generate MSA are described in the Methods section. Those structures *S* differ by the numbers of sequences **A** having large values of *P_nat_*(*S*|**A**), or, informally speaking, by how easy they are to design (Fig. 1). The designability of each structure can be empirically estimated from the diversity between the sequences in the corresponding MSA, from numerical simulations [18], from the properties of the contact map [22], and, as shown in this paper, from the entropy of the inferred Potts model (Methods and SI, Table I).

**Figure 2:**
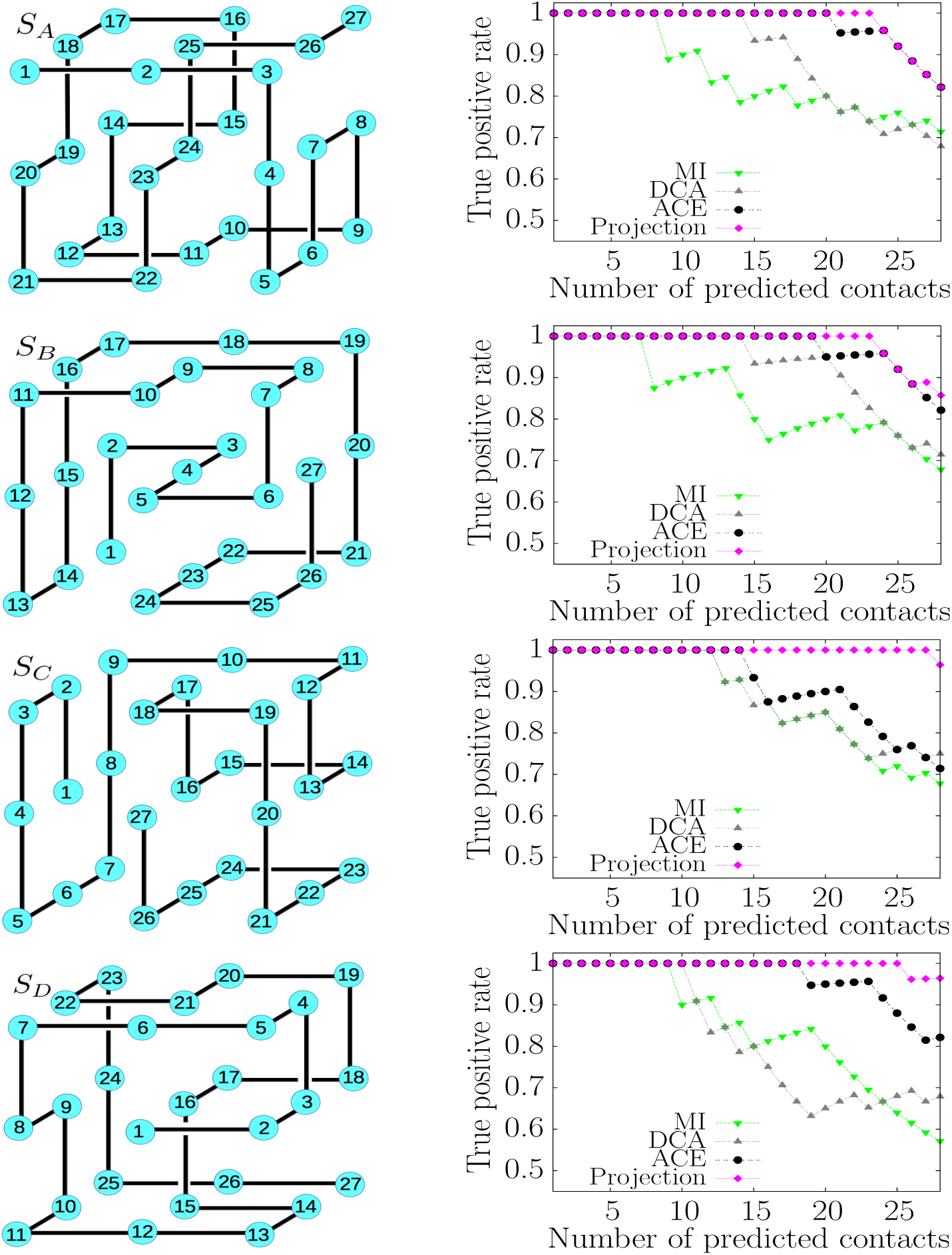
Inverse statistical approaches are able to extract structural information from sequence data of lattice proteins. *Left:* Structures *S_A_*, *S_B_*, *S_C_*, *S_D_*. Amino acids (blue circles) are shown with their number, running from 1 to 27 along the protein backbone (black line). There are 28 contacts between nearest-neighbor amino acids not supported by the backbone. *Right:* True positive (TP) rates, defined as the fraction of contacts among the *k* top scores, with the MI, DCA, ACE, and Projection procedures (Methods). Multi-sequence alignments with *M* = 5 · 10^4^ sequences were generated with Monte Carlo sampling at inverse temperature *β* = 10^3^(Methods).

### Potts pairwise couplings give accurate contact prediction

We show in Fig. 2 the contact predictions for the four structures *S_A_*, *S_B_*, *S_C_*, *S_D_*, based on the ranking of the mutual information (MI) scores and of the inferred Potts couplings, with the mean-field (DCA) and the adaptive cluster expansion (ACE) procedures (Methods). DCA is a fast, approximate method to infer the couplings, while ACE is slower but more accurate (Methods). As in the case of real protein data, Potts-based contact predictions, either with DCA or ACE, generally outperform MI-based predictions. MI, indeed, does not disentangle large indirect correlations mediated by one or more sites from direct correlations due to contacts. ACE couplings are more precise than their DCA counterparts and accordingly, give better contact predictions for all four structures (SI, Table II). The performance of the three methods for contact prediction vary with the native fold (Fig. 2): less designable structures correspond to stronger constraints over (folding) sequences, to larger Potts couplings, and to better contact prediction. Structures *S_A_* and *S_B_* have the smallest designabilities (Fig. 1 and SI, Table 1); contact prediction is extremely good with ACE, while MI works poorly. For those structures, disentangling direct from indirect correlations, as done through the Potts couplings, is therefore very helpful. Conversely, the sequence space corresponding to structures *S_C_* and *S_D_* are less constrained (Fig. 1): the structurally-induced covariation signals are weak, and so are the inferred couplings, especially for structure *S_C_* (SI, Fig. 4). Interestingly, MI outperforms DCA (but not ACE) for structure *S_D_*. MI, contrary to DCA, does not require to invert the correlation matrix, an operation known to behave badly when the (covariation) signal is weak compared to the sampling noise (due to the finite size of the MSA). Improvements for contact prediction (magenta lines in Fig. 2) will result from the detailed interpretation of the Potts couplings below.

### Potts model generates good folding sequences with high probability, in contrast to Independent-site Model

We now test the ability of the inferred Potts model to be generative, that is, to produce sequences having high probability to fold in the native structure. To do so we infer the pairwise Potts model (7) from the MSA, hereafter referred to as ‘natural’, of low-designability structure *S_B_* with the ACE procedure. We sample the Potts distribution using Monte Carlo simulations, thereby generating a new MSA, hereafter referred to as ‘Potts-ACE’. We then compute the probabilities of folding (into *S_B_*) of all the sequences in the Potts-ACE MSA, see Eq. [5]. Results are shown in Fig. 3. Strikingly, the vast majority of the sequences have high folding probabilities. Conversely, sequences generated with an IM following the same procedure are unlikely to have the right fold, as shown in Fig. 3. Pairwise interactions are therefore crucial to design new sequences with the right fold.

**Figure 3:**
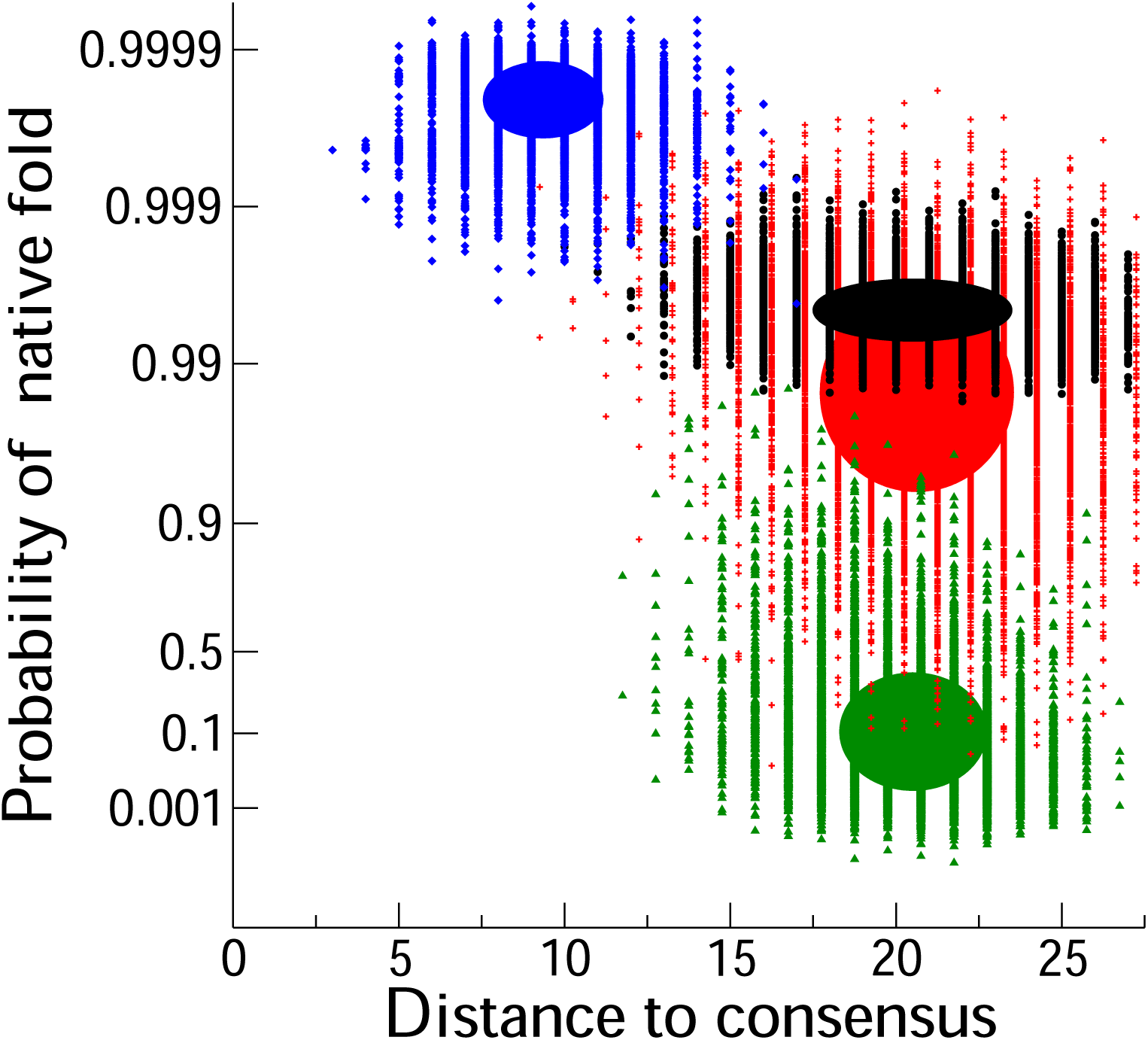
Inferred Potts-ACE model generates sequences with high folding probabilities and diversities. Folding probabilities *P_nat_* (*S_B_* |**A**), Eq. [5], for three sets of 10^4^ sequences **A** randomly generated with the Independent-site Model (IM, green), the Potts-ACE (red) and the Potts-Gaussian (blue) models vs. their Hamming distances to the consensus sequence of the ‘natural’ MSA of structure *S_B_* used to infer the three models. Black symbols show results for the ‘natural’ sequences, sampled with a Monte Carlo procedure (Methods). Hamming distances for the Potts-ACE and IM model have been shifted by, respectively, 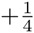 and 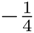 to improve visibility. Filled ellipses show domains corresponding to one standard deviation. Most sequences drawn from the Potts-ACE model have high folding probabilities, while most sequences drawn from the IM have low values of *P_nat_*. Sequences drawn from the Potts-Gaussian model have very high folding probabilities, but are very close to the consensus sequence, and fail to reproduce the diversity of sequences seen in the ‘natural’ MSA (black) and Potts-ACE (red) data.

We stress that Potts couplings have to be inferred with high precision to generate high-quality sequences. We show in Fig. 3 that sequences generated by the Potts-Gaussian model, which makes use of the approximate DCA couplings (details in SI), have very high folding probabilities, but are extremely concentrated around the consensus sequence. The Potts-Gaussian model, contrary to the IM and the Potts-ACE model, fails to reproduce the diversity of sequences observed in the natural MSA. This failure is a direct consequence of the Gaussian approximation (SI, Section V). In summary, the Potts-ACE model shows a drastic improvement over IM and DCA-based models to generate a large set of diverse sequences that fold with high probability.

### Potts couplings reflect both energetics in the native fold and competition with other folds

We now study in more detail the properties of the Potts couplings. We show in Fig. 4A the scatter plot of the inferred couplings *J_ij_* (*a, b*) for structure *S_B_* vs. the Miyazawa-Jernigan energetics parameters − *E*(*a, b*) used to compute the LP energies (Methods); scatter plots for the other structures are shown in SI. For each pair of sites *i, j*, we observe a linear dependency,

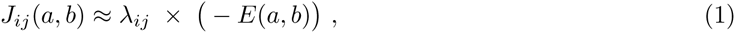

where the slope *λ_ij_* is positive for the pairs in contact in the native structure (red symbols in Fig. 4A), and negative, or zero for the pairs not in contact (blue symbols). In the following, we characterize each pair *i, j* with its slope, computed through

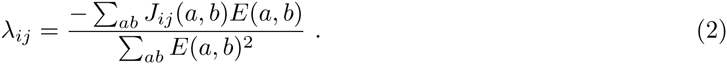

**Figure 4:**
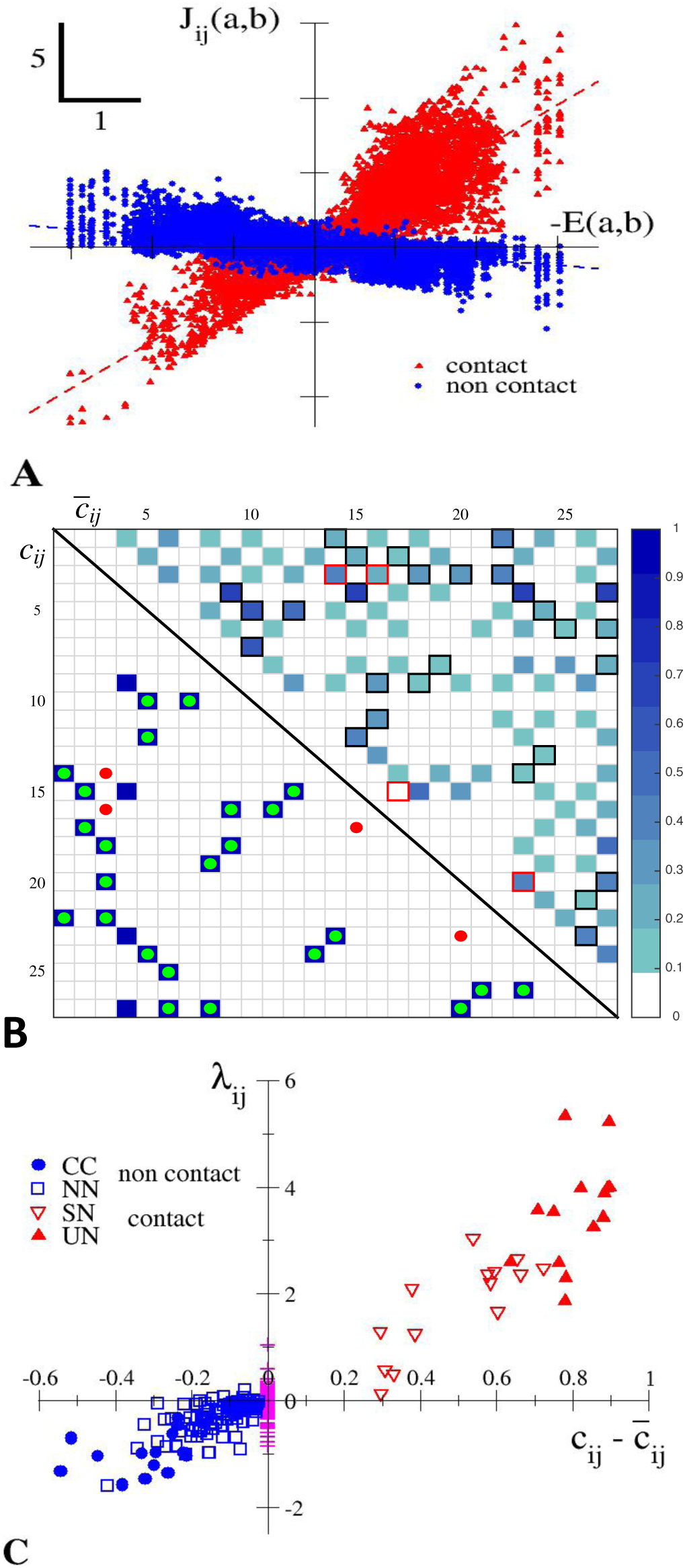
Inferred Potts couplings encode energetics and structural information about native and competitor folds, reflecting both positive and negative designs. **A.** Values of *J_ij_* (*a, b*) (inferred from a MSA of structure *S_B_*) vs. −*E*(*a, b*) across all pairs of sites *i, j* and of amino acids *a, b* (found at least once in the MSA on those sites). Red symbols correspond to pairs (*i, j*) in contact, while blue symbols correspond to no contact. Broken lines: linear fits for the pairs in contact (red, slope ⋍ 3.16) and not in contact (blue, slope ⋍ −0.41). **B.** Lower-triangle: contact map *c_ij_* of structure *S_B_*. Full blue squares: pairs of sites *i, j* in (green dots) and not in (red dots) contact among the 28 largest scores 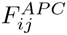 (Methods). Upper triangle: average contact map 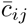, computed over all competitor folds weighted with their Boltzmann weights (SI). Contacts not present among the 28 top scores, and three out of four false positive predictions (red dots in the lower triangle, shown with red squares) correspond to large 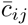; four of them are carried by the center of the cube (*i* = 4 on *S_B_* and *S_F_*, see Fig. 2). **C.** Pressure *λ_ij_* for each pair of sites (*i, j*), computed from Eq. [2], vs. 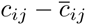 for structure *S_B_*. The 28 contacts on *S_B_* (red symbols) are partitioned into the Unique-Native (UN, 14 full up triangles) and Shared-Native (SN, 14 down triangles) classes, according to, respectively, their presence or absence in the closest competitor structure, *S_F_*. The 128 non-contact pairs on *S_B_* (blue symbols) are partitioned into the Closest-Competitor (CC, 14 full circles) and the Non-Native (NN, 114 squares) classes, with the same criterion as for contacts. The 195 pairs of sites which can never be in contact on any fold due to the lattice geometry are shown with magenta pluses. Theoretical predictions from Eq. [3], averaged over the pairs of sites in the UN, SN and CC classes, give 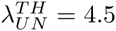, 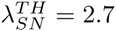 and 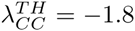, in good agreement with the average values of the slopes: *λ_UN_* ≈ 4 ± 1.5, *λ_SN_* ≈ 2 ± 1.5, *λ_CC_* ≈ −1 ± 0.5. Similar behaviours are found for *S_A_*, *S_C_* and *S_D_*, see SI, Table III and Figs. 7, 8 and 9.

We interpret *λ_ij_* as a measure of the coevolutionary pressure on the sites *i*, *j*, due to the design of the native structure. This interpretation is supported by the following theoretical and approximate expression for the pressure (see Methods and derivation in SI):

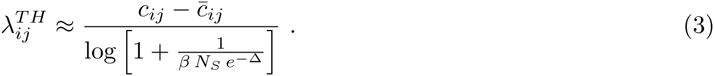

The numerator 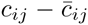 measures the variation between the native contact map *c_ij_* (= 1 if *i, j* are in contact and 0 otherwise), and the average contact map 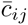 of the structures in competition with the native fold, each weighted by *e*^−Δ^, where Δ is a the typical gap between the energy of sequences folded into the native structure and their energies in the competing structures (Methods, Eq. [9] and below) [23]. The native and average contact maps for structure *S_B_* are shown, respectively, in the lower and upper triangles of Fig. 4B. The denominator in Eq. [3] depends on the inverse sampling temperature *β* in the Monte Carlo procedure to generate the MSA, the effective number *N_S_* of competing structures, and the typical energy gap Δ (Methods).

Figure 4C shows the pressures *λ_ij_* computed from the inferred couplings, Eq. [2], vs. the contact-map difference 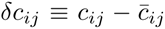 across all pairs of sites *i, j* for the native fold *S_B_*. We observe a monotonic dependence of the pressure *λ_ij_* with *δc_ij_*; in particular, *λ_ij_* has the same sign as *δc_ij_*. Note that pairs *i, j* such that *j* – *i* is even valued can never be in contact in any structure on a cubic lattice: 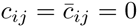, as can be seen in the checkerboard pattern of 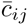 in Fig. 4B. The associated pressures *λ_ij_* are weak and indicate the level of noise in the inference (magenta symbols in Fig. 4C).

Pairs with large and positive *δc_ij_* are in contact in the native fold, but not in the competitor structures. They are under strong covariation pressure to stabilize the native fold and not the competitors (positive design). On the contrary, pairs with small and positive *δc_ij_* are in contact both in the native and competitor folds, and are subject to weak pressures. As a result, couplings are weak, and contact prediction is difficult. For the structures we have studied, those true negatives generally correspond to the contacts between the site at the center of the cube and its four neighbors, as these contacts are also shared with the closest competitor folds (Fig. 4B).

Sites *i, j* not in contact in the native fold correspond to negative *δc_ij_*. Most of those pairs are neither in contact in the competitor structures, and correspond to weak pressures and inferred couplings. However, some pairs do correspond to contact in the competitor folds (large and negative *δc_ij_*), and are subject to strong and negative pressures. The anti-correlation between the couplings *J_ij_* (*a*, *b*) and the Myiazawa-Jernigan parameters *E*(*a*, *b*) is the result of negative design: the native structure is favored by rendering contacts that appear only in competitor folds unfavorable [24].

A detailed classification of the pairs of sites is shown in Fig. 4C, based on the identification of the closest competitor to *S_B_*, structure *S_F_* (SI, Fig. 10). It allows us to define four classes of contacts, and to illustrate concretely the positive and negative design mechanisms elucidated above [21,24].

The pressures *λ_ij_* can be used for contact prediction, and give much better TP rates than classical estimators based on the norm of the couplings, *F_ij_* in Eq. [8], see ‘projection’ TP rates in magenta in Fig. 2. Large *F_ij_* not corresponding to contacts can be either due to sampling noise in the inferred couplings, incoherent with *E*(*a*, *b*), or to large couplings anti-correlated with *E*(*a, b*); the latter give rise to negative *λ_ij_*, and the former to small *λ_ij_*.

As the scores used for contact predictions are based on the squared couplings (Methods), negative pressures give rise to large scores, and to false positives.

To further test the theoretical expression of the pressure, Eq. [3], we have varied two features of the Monte Carlo procedure used to generate the MSA: (1) the pool of possible competing structures appearing in the denominator of the folding probability, *P****_nat_*** in [5], and (2) the inverse sampling temperature *β* (Methods). We observe that the pressures *λ_ij_* increase when the pool of competing structures is restricted to structures similar to the native one, or when *β* is increased, as sequences are constrained to have higher values for *P_nat_* (SI, Fig. 5 and Table III).

## Discussion

Lattice proteins offer a fully controlled, non–trivial benchmark to understand the factors that affect the success of inverse statistical approaches for structural prediction from multi sequence alignments (MSA). Structural prediction based on the maximum entropy Potts model inferred to reproduce the low-order statistics of MSA outperforms Mutual Information approaches, as is the case for real protein data, especially when couplings are accurately inferred (ACE procedure) (Fig. 2). An important finding in the present work is the approximate linear relationship between the inferred Potts couplings *J_ij_*(*a*, *b*) and the Miyazawa-Jernigan energetic parameters *E*(*a, b*) across the pairs of amino acids *a, b* for a given pair of sites *i, j*, Eq. [1] and Fig. 4A. The slope of this linear relationship, *λ_ij_*, is a measure of the covariation ‘pressure’ on the pair *i, j* due to the structural constraints. The pressure depends on *(i)* the difference between the contact maps of the native fold (*c_ij_*) and the ones of the competing structures 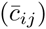, see Fig. 4B; *(ii)* the proximity (in terms of energy, Eq. [9]) of the native fold with its competitor structures; *(iii)* the strength of constraints in the design, controlled here by the inverse sampling temperature (Eq. [3]). A consequence of *(i)* is that the intensity of the inferred couplings *J_ij_*(*a, b*), measured through the scores *F_ij_* in Eq. [8], are directly related to the local properties of the contact map on sites *i, j* (and not on other sites), which explains the success of inverse approaches to disentangle direct from indirect effects in amino-acid correlations. However, the relationship also involves the average contact map 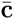, originating from the competitor folds. As 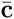, contrary to the native contact map **c**, is not sparse, so are the couplings **J**. Couplings can, in particular, take non-zero values for *i, j* in contact one the competing folds (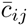 close to 1), but not in the native fold (*c_ij_* = 0), see Figs. 4A,B. Those ‘repulsive’ couplings give rise to negative pressures *λ_ij_*, and prevent the sequence from forming contacts that would result in the wrong fold, a clear illustration of negative design [21,24,25]. Reciprocally, sites in contact in the native fold (*c_ij_* = 1) need not be subject to strong covariation and associated to strong couplings, if they are also in contact in the competing folds (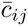 close to 1). Our results clearly show that strong couplings do not necessarily imply nor require structural contact (in the native fold). They also explain the success and limitations of contact prediction based on the magnitude of the inferred couplings (Fig. 2).

A second major result of our approach, shown in Fig. 3, is that the inferred Potts model, when accurately inferred, here, with the ACE procedure, generates sequences that fold with high probability in the native state and show amino-acid diversity similar to the one observed in the original MSA. This performance is remarkable. While the energy of a LP sequence in a given structure is a sum of pairwise contributions only, the effective Hamiltonian of the sequences constrained to fold with high probability in this structure (and not in the competing folds) includes high-order multi-body interactions (originating from the competing folds) between amino acids, as can be expected for real proteins. Furthermore, independent-site models are totally unable to produce sequences with the correct fold (Fig. 3). Our result corroborates the works of Ranganathan and collaborators [13,14], who experimentally showed that sequences built according to a reshuffling of the MSA of the small WW domain respecting site conservation and (approximately) pairwise correlations folded correctly with a good (30%) probability. The generative character of the Potts model, combined with the linear relationship between couplings and energetics parameters, agrees with previous studies showing that designed and real protein-like sequences are generated by Gibbs measure with Hamiltonian reflecting real protein energetics [17,24]. We stress that couplings have to be inferred with great care, going beyond the mean-field DCA approximation, in order for the Potts model to be generative (Fig. 3).

The analysis of real protein data could benefit from our analysis in several ways. First, the understanding of the relationship between the couplings and the contact map obtained here indicates that the pressures *λ_ij_* (Eq. [2]) are excellent estimators of contacts (Fig. 2), outperforming the usual scores 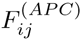 (Eq. [8]). The use of *λ_ij_* allows one to both keep track of the sign of the couplings (and avoid false positives corresponding to pressures originating from negative design, see above) and increase signal-to-noise ratio, by removing noise in the inferred couplings not aligned along the Myiazawa-Jernigan energy matrix. In real proteins, the physico-chemical properties of amino acids suggest the existence of difference classes of contacts, whose energetics could be inferred from large-scale analysis of databases and used to define class templates E. Projections of the inferred Potts couplings on those templates could in turn be used for contact predictions. Secondly, our work suggests a practical way to quantify the designability of a protein family, *i.e*. to measure the number of sequences ‘belonging’ to the family. Current approaches rely on the computation of the maximal eigenvalue of the contact matrix **c** [22], though our study suggests that the matrix 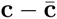 should rather be considered. Given the generative character of the inferred Potts model, a direct and more accurate measure of the designability is provided by its entropy computed with the ACE procedure, see Methods and SI, Table I. This approach can be applied to real protein families, an example is provided in [26].

In future studies, the total control we have over lattice proteins will allow us to study the importance of covariation and the success of inverse approaches in ’Gedanken’ experiments related to many important issues of interest for real proteins, and going beyond the structural aspects studied here. Among those let us cite the detection of homology. Current protein databases, such as PFAM, classify query sequences into families based on Hidden Markov Model profiles, an extension of independent-site models capable of taking into account deletions and insertions [27]. It is an open problem to understand whether coupling-based models, exploiting covariation, could, contrary to HMM, recognize sequences with low homology in a reliable and computationally tractable way. Another very important issue is the estimation of fitness landscapes, or fitness changes in responses to one or more mutations [28, 29]. Covariation-based models have been recently introduced to predict escape paths of pathogens (virus or bacteria) from drugs or vaccines in this context [15]. Lattice proteins offer a unique benchmark to understand deeply and quantitatively and, ultimately, improve those approaches.

## Materials and Methods

### Lattice-protein model

We consider model proteins, whose *L* = 27 amino acids occupy the sites of a 3 × 3 × 3 cubic lattice [16-18]. Four of the 103, 406 possible configurations of the backbone (excluding symmetries), hereafter called folds or structures, are shown in Fig. 2. Unless otherwise said, we restrict ourselves to a representative subset of 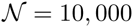 folds [30]. Two amino acids are said to be in contact if they are nearest neighbors on the lattice (but not on the backbone). The contact map **c^(^*^S^*^)^** of structure *S* is its 27 × 27 adjacency matrix: 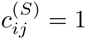 if *i, j* are in contact, 0 otherwise. Amino acids in contact interact through the Miyazawa-Jernigan (MJ) 20 × 20 energy matrix *E* [19]. The energy of a sequence **A** = (*a***_1_**, &, *a***_27_**) of amino acids folded into structure *S* is

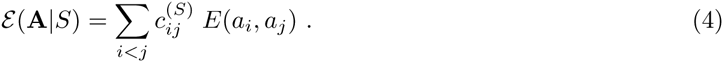

The Boltzmann probability that sequence **A** folds into structure *S* is given by (for unit temperature):

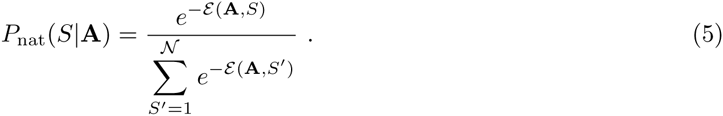

Good folders are sequences **A** with large gaps between 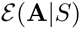 and the energies 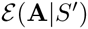 with the other folds [17].

### Sampling of sequence space

We generate a multi-sequence alignment (MSA) for the native fold, say, *S*, through Monte Carlo simulations with the Metropolis rule [24]. The simulation starts from a sequence **A**, and attempts to mutate randomly one amino acid at a time, say, 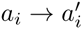; Let **A**′ denote the new sequence. If *P***_nat_**(*S*|**A**′) > *P***_nat_**(*S*|**A**) the mutation is accepted; otherwise it is accepted with probability [*P***_nat_**(*S*|**A**′)/*P***_nat_**(*S*|**A**)]*^β^*(< 1). Parameter *β* plays the role of an inverse algorithmic temperature that sets the stringency of sequence selection in the sampling Monte Carlo procedure. In practice we choose *β* = 10**^3^**, which ensures that sequences fold in the native structure S with probability 0.995 or larger and that thermalization is fast (SI, Fig. 1). The MSA is made of the sequences **A***^τ^*, with *τ* = 1, &, *M* ~ 10**^4^** − 10**^5^**, generated at regular intervals of time, larger than the correlation time of the Monte Carlo dynamics. Each sequence **A***^τ^* is therefore drawn according to the equilibrium measure equal to *P*_nat_(*S*|**A***^τ^*)*^β^*, up to a normalisation multiplicative factor. The corresponding effective Hamiltonian includes not only the native-structure energy 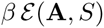 in Eq. [4], with pairwise interactions between the amino acids *a_i_*, but also contributions coming from all the other folds *S*′, with multi-body interactions at any orders ≥ 2.

### Inference of Potts model from sequence data

We consider the maximum entropy (least constrained [8]) distribution over the sequences, that reproduces the one- and two-point amino-acid empirical frequencies computed from the sequences **A***^τ^* in the MSA of a given native structure,

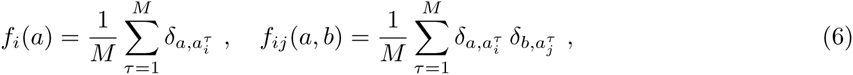

where *δ* denotes the Kronecker function. This maximum entropy distribution is the Gibbs measure of a

Potts model, with *q* = 20-state variables, whose energy is given by

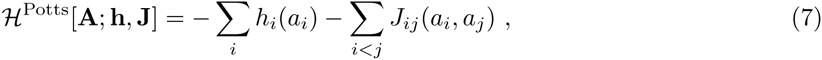

where the sets of *L* × *q* fields, **h** = {*h_i_*(*a*)}, and of *L*(*L* − 1)/2 × *q***^2^** couplings, **J** = {*J****_i_****_j_*(*a*, *b*)}, are chosen so that the frequencies computed from the Potts model distribution reproduce those of the data, Eq. [6].

We resort to two methods to solve this hard computational problem. Within the mean-field Direct Coupling Approximation (DCA), already applied to many real protein data [6], **J** is approximated as minus the pseudo-inverse of the connected correlation matrix **c**, with entries *c_ij_*(*a, b*) = *f_ij_*(*a, b*) − *f_i_*(*a*)*f_j_*(*b*). The Adaptive Cluster Expansion (ACE) is a more accurate but slower procedure, which recursively builds the most relevant clusters of strongly interacting sites [31,32]. In addition to **h**, **J** ACE gives access to the entropy of the inferred Potts model, which is an estimate of the designability of the native fold. See SI, Section II for details about the implementation of ACE, and the ability of the Potts-ACE distribution to reproduce high-order statistics of the MSA.

For the Independent-site Model (IM), the energy 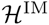 is a sum of field contributions only (**J** = 0 in Eq. [7]); the corresponding fields are simply given by 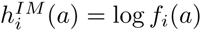.

### Contact prediction

Computation of the Potts couplings **J** allows us to define the scores [6] (with the Averaged Product Correction [33])

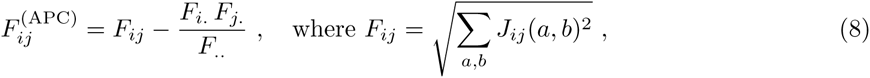

and *F_i_*. = ∑*_l_ F_il_*, *F*. = ∑*_i_ F_i._*. Scores are then ranked in decreasing order, and used to predict contacts. The true positive rate at rank *k* is the fraction of pairs of amino acids (*i, j*) in contact among the top *k* scores 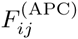. Within the mutual information approach, scores are simply given by 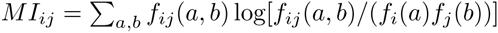 [34], and do not distinguish direct from indirect correlations.

### Competitors, gap and average contact map

We define the gap Δ(*S*|*S_nat_*) between structure *S* and the native fold *S_nat_* through

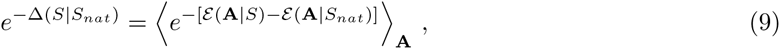

where the average is taken over sequences **A** in a MSA of *S_nat_*. The distributions of gaps for *S_nat_* = *S_A_*, *S_B_*, *S_C_*, *S_D_* are shown in SI, Fig. 6. We approximate the number *N_S_* of competitor folds and their typical gap Δ with the native structure *S_nat_* through 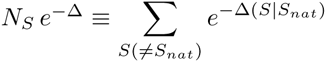. The average contact map of those competitors is defined as 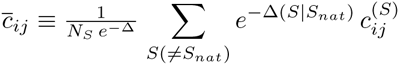.

## Acknowledgments

We are grateful to M. Weigt for many useful discussions. H.J., S.C. and R.M. are funded by ANR-13-BS04-0012-01 (Coevstat). A.G. is funded by NSF predoctoral fellowship and E.S. is funded by NIH RO1 068670.

## Supporting Information Files

**Text S1.** Supporting Information for Benchmarking inverse statistical approaches for protein structure and design with exactly solvable models.

